# Perineuronal net receptor PTPσ regulates retention of memories

**DOI:** 10.1101/2021.02.26.433000

**Authors:** Angelina Lesnikova, Plinio Casarotto, Caroline Biojone, Eero Castrén

**Author notes:** Corresponding authors: Caroline Biojone, Eero Castrén.

## Abstract

Perineuronal nets (PNNs) have an important physiological role in retention of learning by restricting cognitive flexibility. Their deposition peaks after developmental periods of intensive learning, usually in late childhood, and they help in long-term preservation of new acquired skills and information. Modulation of PNN function by various techniques enhances plasticity and regulates retention of memories, which may be beneficial when memory persistence entails negative symptoms such as post-traumatic stress disorder (PTSD). In this study, we investigated the role of PTPσ (receptor-type tyrosine-protein phosphatase S, a phosphatase that is activated by binding of chondroitin sulfate proteoglycans from PNNs) in retention of memories using novel object recognition and fear conditioning rodent models. We observed that mice haploinsufficient for *PTPRS* gene (PTPσ^+/−^), although having improved short-term object recognition memory, display impaired long-term memory in both novel object recognition and fear conditioning paradigm, as compared to WT littermates. However, PTPσ^+/−^ mice didn’t show any differences in behavioral tests that do not heavily rely on cognitive flexibility, such as elevated plus maze, open field, marble burying and forced swimming test. Since PTPσ has been shown to interact with and dephosphorylate TRKB, we investigated activation of this receptor and its downstream pathways in limbic areas known to be associated with memory. We found that phosphorylation of TRKB and PLCγ are increased in the hippocampus, prefrontal cortex and amygdala of PTPσ^+/−^ mice, but other TRKB-mediated signaling pathways are not affected. Our data suggest that disruption of PNN-PTPσ complex facilitates short-term memory by promoting TRKB phosphorylation in different brain areas, but that PTPσ activity is required for the retention of long-term memories. Inhibition of PTPσ or disruption of PNN-PTPσ-TRKB complex might be a potential target for disorders where negative modulation of the acquired memories can be beneficial.

**Conflict of interest:** The authors declare no conflict of interest.

**Funding:** This work was supported by Doctoral Program in Integrative Life Science, Jalmari ja Rauha Ahokkaan Säätiö grant, Centre for International Mobility (CIMO) Grant TM-16-10112, and by grants from European Research Council (#322742), EU Joint Programme - Neurodegenerative Disease Research (JPND) CircProt (#301225 and #643417), Sigrid Jusélius Foundation, Jane and Aatos Erkko Foundation, and the Academy of Finland (#294710 and #307416). None of the funders had a role in the data acquisition, analysis or manuscript preparation.

**Author contributions:** PC, CB and EC designed the study; AL, PC and CB performed the experiments and analyzed the data; AL wrote the manuscript draft; AL, PC, CB and EC revised the manuscript.

**Contribution to the Field Statement:** Plasticity of neuronal networks increases brain’s ability to adapt and it is compromised in various conditions including psychiatric diseases, brain injuries, neurodevelopmental and neurodegenerative disorders. Perineuronal nets, through their receptor PTPσ restrict neuronal plasticity in adult brain, which is considered important for the retention of long-term memories. We have previously shown that this restricted plasticity is mediated by inhibition by PTPσ of the TRKB neurotrophin receptor. We have now investigated the biochemical and behavioral phenotype of mice with reduced PTPσ expression. We observed that PTPσ^+/−^ mice have increased phosphorylation of TRKB and its downstream partner PLCγ1 in the prefrontal cortex, hippocampus and amygdala, demonstrating chronic overactivation of plasticity-related pathways. Consistently, PTPσ^+/−^ mice demonstrated facilitated learning but impaired long-term memory retention. Unexpectedly, long-term memory of PTPσ^+/−^ mice was impaired, suggesting that perneuronal net-stimulated PTPσ activity is important in memory retention. This effect of neuronal “hyperplasticity” induced by PTPσ knockout may be both beneficial and harmful in the treatment of human patients, which should be taken into account when developing therapeutic strategies.

## Introduction

Perineuronal nets (PNNs) are complex extracellular matrix structures consisting of a core protein and multiple glycosaminoglycan chains, such as chondroitin sulfate proteoglycans (CSPGs) and heparan sulfate proteoglycans (HSPGs) (Deepa et al., 2006; Testa et al., 2019). They surround somata and proximal dendrites of certain types of neurons (Hockfield and McKay, 1983), being particularly enriched around parvalbumin-positive interneurons (PV+) (Kosaka and Heizmann, 1989; Fawcett et al., 2019). Perineuronal nets are well-recognized regulators of plasticity in the CNS which stabilize and consolidate the system (Banerjee et al., 2017; Christensen et al., 2021), and play an indispensable role in learning and memory (Wang and Fawcett, 2012; Sorg et al., 2016; Shen, 2018). Interference with perineuronal net formation by chondroitinase ABC (chABC) digestion of their sugar chains or genetic knockdown of PNN components has been shown to promote plasticity in multiple paradigms, including ocular dominance (Pizzorusso et al., 2002), drug-induced conditioned place preference (Xue et al., 2014; Slaker et al., 2015) and fear learning (Gogolla et al., 2009; Banerjee et al., 2017; Pignataro et al., 2017; Shi et al., 2019).

The most abundant component of PNNs, CSPGs, have been discovered to exert their inhibitory action on neuronal plasticity by binding to protein tyrosine phosphatase sigma (PTPσ), a transmembrane receptor (Shen et al., 2009; Coles et al., 2011). We have recently found that the PNN-PTPσ complex exerts its inhibitory action on neuronal plasticity by restricting signaling of neurotrophic receptor tyrosine kinase 2 (TRKB) (Lesnikova et al., 2021). TRKB is a receptor for brain-derived neurotrophic factor (BDNF) and a well-known activator of plasticity in the brain (Minichiello et al., 1999; Castrén and Antila, 2017; Umemori et al., 2018). We found that genetic deficiency of PTPσ increases phosphorylation of TRKB and promotes cortical plasticity at network level. We also showed that chABC-induced plasticity in the adult brain requires intact TRKB expression in PV+ interneurons, a neuronal class whose mode of function is particularly dependent on PNNs. There, we propose a model where, under normal conditions in the adult brain, perineuronal nets bind to PTPσ and promote dephosphorylation of TRKB and restriction of its activity, while reduction of interaction between PNNs and PTPσ or PTPσ and TRKB enhances the ability of BDNF to phosphorylate TRKB and promote plasticity.

Here, we set out to further investigate the effects of PTPσ downregulation on neuronal plasticity. In particular, our aim was to characterize the molecular profile of enhanced plasticity induced by PTPσ genetic deficiency, and to reveal any changes in behavioural phenotype of animals haploinsufficient for *PTPRS* gene.

## Materials and Methods

### Animals

Adult BALB/c mice heterozygous for *PTPRS* gene (PTPσ^+/−^) mice and their wild-type (WT) littermates of both sexes were used for the experiments. The animals were group-housed (2-5 animals per cage in type ll: 552 cm2 floor area, Tecniplast, Italy) under standard laboratory conditions with a 12-hour light/dark cycle and access to food and water *ad libitum*. Behavioral experiments were carried out during the light phase of the cycle. All the procedures involving animals were carried out with approval of the Experimental Animal Ethical Committee of Southern Finland (ESAVI/38503/2019).

### Antibodies

The following antibodies were used in the study: anti-Akt rabbit polyclonal antibody (pAb) (Cell Signaling Technology, #9272); anti-β-Actin mouse monoclonal antibody (mAb) (Sigma-Aldrich, #A1978); anti-p44/42 MAPK (Erk1/2) rabbit pAb (Cell Signaling Technology, #9102); anti-p70 S6 kinase rabbit pAb (Cell Signaling Technology, #9202); anti-phospho-Akt rabbit mAb (Cell Signaling Technology, #4056); anti-phospho-p44/42 MAPK (Erk1/2) rabbit mAb (Cell Signaling Technology, #9101); anti-phospho-p70 S6 kinase rabbit polyclonal antibody (pAb) (Cell Signaling Technology, #9204); anti-phospho-PLCγ1 rabbit pAb (Cell Signaling Technology, #2821); anti-PLCγ1 rabbit mAb (Cell Signaling Technology, #5690); anti-phospho-TRKA (Tyr490)/TRKB (Tyr516) rabbit mAb (Cell Signaling Technology, #4619); anti-PSD-93/anti-Chapsyn-110 mouse mAb (UC Davis/NIH NeuroMab Facility; anti-PSD-95 mouse mAb (UC Davis/NIH NeuroMab Facility); anti-TRKB goat pAb (R&D Systems, #AF1494); secondary goat anti-mouse horseradish peroxidase (HRP)-conjugated antibody (Bio-Rad, #1705047); secondary goat anti-rabbit HRP-conjugated antibody (Bio-Rad, #1705046), secondary rabbit anti-goat HRP-conjugated antibody (Invitrogen, #611620).

### Brain sample collection and processing

The animals were euthanized by CO_2_. The prefrontal cortex, hippocampus and amygdaloid complex were dissected and immediately placed on dry ice. The samples were homogenized by sonication in NP lysis buffer (137mM NaCl, 20mM Tris, 1% NP-40, 10% glycerol, 48mM NaF) containing a protease and phosphatase inhibitor mix (#P2714 and #P0044, Sigma Aldrich) and 2mM Na_2_VO_3_. The homogenized samples were centrifuged for 15 min at 15000 G, 4° C. The supernatant was collected and stored at −80° C until further use. Protein quantification was done by colorimetric Lowry method using DC Protein Assay Kit (Bio-Rad, #5000116) and the product of the colorimetric reaction read at 750 nm in a Varioskan Flash plate reader (Thermo Fisher Scientific).

### Enzyme-linked immunosorbent assay (ELISA)

Samples from female mice 2-3 months old were used for this experiment. We assessed levels of TRKB phosphorylation (pTRKB) using a protocol previously described by Antila et al. (Antila et al., 2014). A high-binding 96-well OptiPlate (Perkin Elmer) was coated with anti-TRKB antibody incubated in carbonate buffer (57.4 mM sodium bicarbonate and 42.6 mM sodium carbonate, pH 9.8) 1:500 at 4° C overnight. On the next day, the antibody was removed, and the non-specific binding was blocked using a blocking buffer containing 3% bovine serum albumin (BSA) (Sigma-Aldrich, #A9647) in PBST (137 mM NaCl; 10 mM Phosphate; 2.7 mM KCl; 0.1% Tween-20; pH 7.4) for 2 hours. After that, homogenized and centrifuged brain lysates (80 μg of total protein) were transferred to the plate and incubated at 4° C overnight. On day 3, the brain lysates were removed, the plate was washed 4 times with PBST using a plate washer (Thermo Fisher Scientific Wellwash Versa), and anti-pTRKB antibody (1:1000 in the blocking buffer) was incubated at 4° C overnight. On the final day, the antibody was removed, the plate was washed 4 times with PBST, and incubated with horseradish peroxidase (HRP)-conjugated antibody (1:5000 in the blocking buffer) for 1 hour at room temperature. The antibody was removed, the plate was washed 4 times with PBST, and HRP ECL substrate (Thermo Fisher Scientific, #32209) was incubated in the plate for 3 min. Chemiluminescence was measured using 1 s integration time in a Varioskan Flash plate reader (Thermo Fisher Scientific).

### Western blot

Samples from male and female mice 2 months old were used for this experiment. 40 μg of total proteins from each sample was heated in 2X Laemmli buffer (4% SDS, 20% glycerol, 10% 2-mercaptoethanol, 0.02% bromophenol blue and 125mM Tris HCl, pH 6.8) for 5 min at 95° C and loaded to NuPAGE 4-12% Bis-Tris Protein polyacrylamide gels (Invitrogen, #NP0323BOX). After electrophoresis, the proteins were transferred to a polyvinylidene difluoride (PVDF) membrane, blocked in the blocking buffer (3% BSA in Tris-phosphate buffer: TBST, 20 mM Tris–HCl; 150 mM NaCl; 0.1% Tween-20; pH 7.6) and incubated with the primary antibody diluted 1:1000 in the blocking buffer overnight at 4° C. The membranes were subsequently incubated in HRP-conjugated secondary antibodies diluted in the blocking buffer (1:10000) for 1 hour at room temperature. The signal was detected using enhanced chemiluminescent (ECL) HRP substrate WesternBright Quantum (Advansta, #K-12042) in a CCD camera (G:Box Chemi, Syngene). The levels of phospho-proteins were normalized by total levels of the corresponding proteins; the levels of total TRKB, PSD-93 and PSD-95 were normalized by B-actin. PSD-95 samples from PFC were not normalized due to low detection signal from the housekeeping protein.

### Novel Object Recognition test

Male and female mice 3-4 months old were used for these experiments. NOR test was performed in a transparent arena (39×20×16 cm) where two identical objects (white plastic ping-pong balls glued to 50-ml Falcon tube caps) holders at the bottom) were located (adapted from (Casarotto et al., 2021)). The animals were allowed to explore the objects for 15 min in 3 consecutive days (training sessions). At the test session (4 hours or 5 days after the last training session to assess short-term or long-term memory, respectively) one of the old objects was replaced by a new one (black rubber squash ball glued to 50-ml Falcon tube caps), and the animals were allowed to explore both objects for 5 min. The number of interactions with the objects was calculated, and the difference between the visits to the new object and the old object (fB-fA) was used to assess memory. Total number of visits (fA+fB) was used to assess total exploratory activity. Two different cohorts of animals were used to assess short-term and long-term memory.

### Fear Conditioning

Male mice 4-7 months old were used for this experiment. Mice were subjected to fear conditioning in a squared arena A (23 cm side, 35 cm height) with a metal grid on the floor and transparent acrylic glass walls, with continuous background white noise and 100 lux illumination. On day 1, the animals received five foot shocks (0.6 mA, 1 s) each preceded by a 30-s sound cue (latency to the first foot shock: 150 s, time between the last foot shock and the end of the trial: 30 s, inter-shock interval: 30-120 s, total session length: 8 min 45 s). On day 10, the mice were first placed in a new arena B (same size as arena A but with plastic grey floor and plastic black walls), and freezing in response to the sound cue was measured (cue retrieval). Two hours later, the mice were placed in the same context where fear conditioning took place (arena A), and freezing in response to the sound cue was measured (context + cue retrieval). Behavior was counted as freezing if the mouse was not moving for at least 3 s, and measurements were done automatically by TSE system software.

### Elevated Plus Maze

Male mice 3 months old were used for this experiment. The apparatus consisted of a central zone (5 × 5 cm), two open arms (30 × 5 cm) and two closed arms (30 × 5 cm) surrounded by transparent acrylic glass walls (15 cm height). The maze was elevated 40 cm above the floor. The illumination was set to ~150 lux. The animal was placed in the center of the arena and allowed to explore the maze for 5 min. The animal activity was tracked automatically by the Noldus EthoVision XT 8 system.

### Open Field test

Male mice 3 months old were used for this experiment. The test was carried out in a squared arena (30 cm side) with transparent walls (20 cm) and white floor. The illumination was set at ~100 lux. The mice were placed in the center of the arena and allowed to explore it freely for 5 min. The activity was tracked automatically by Noldus EthoVision XT 8 system.

### Marble Burying test

Male mice 3 months old were used for this experiment. 12 glass marbles were evenly distributed on a fresh 5 cm-deep wood chip bedding in a rectangular plastic arena similar in size to the mouse home cage (39×20×16 cm). The mice were given 15 min to explore the marbles, and the number of buried marbles was counted. A marble was considered buried if at least 2/3 of its surface was covered with bedding chips (Njung’e and Handley, 1991).

### Forced Swimming test

Male mice 3 months old were used for this experiment. The animals were placed in a 5-liter 19-cm ø glass cylinder with a 20-cm water column at room temperature (~22° C) and were left there for 6 min. Percentage of time spent floating during the last 4 min (immobility time) was used for the analysis.

### Statistical analysis

The data were analyzed using Graph Pad Prism 6 software. Parametric tests were preferentially used when possible, non-parametric tests were used when the values were discrete or homoscedasticity was not observed. ROUT test was used to identify outliers (Motulsky and Brown, 2006). Detailed information on statistical tests and values is provided in Table 1. Differences between groups were considered statistically significant when p < 0.05. Data in the figures are presented as mean ± SEM. All data used in the present study are available under CC-BY license DOI: 10.6084/m9.figshare.c.5316767.

**Table 1.**
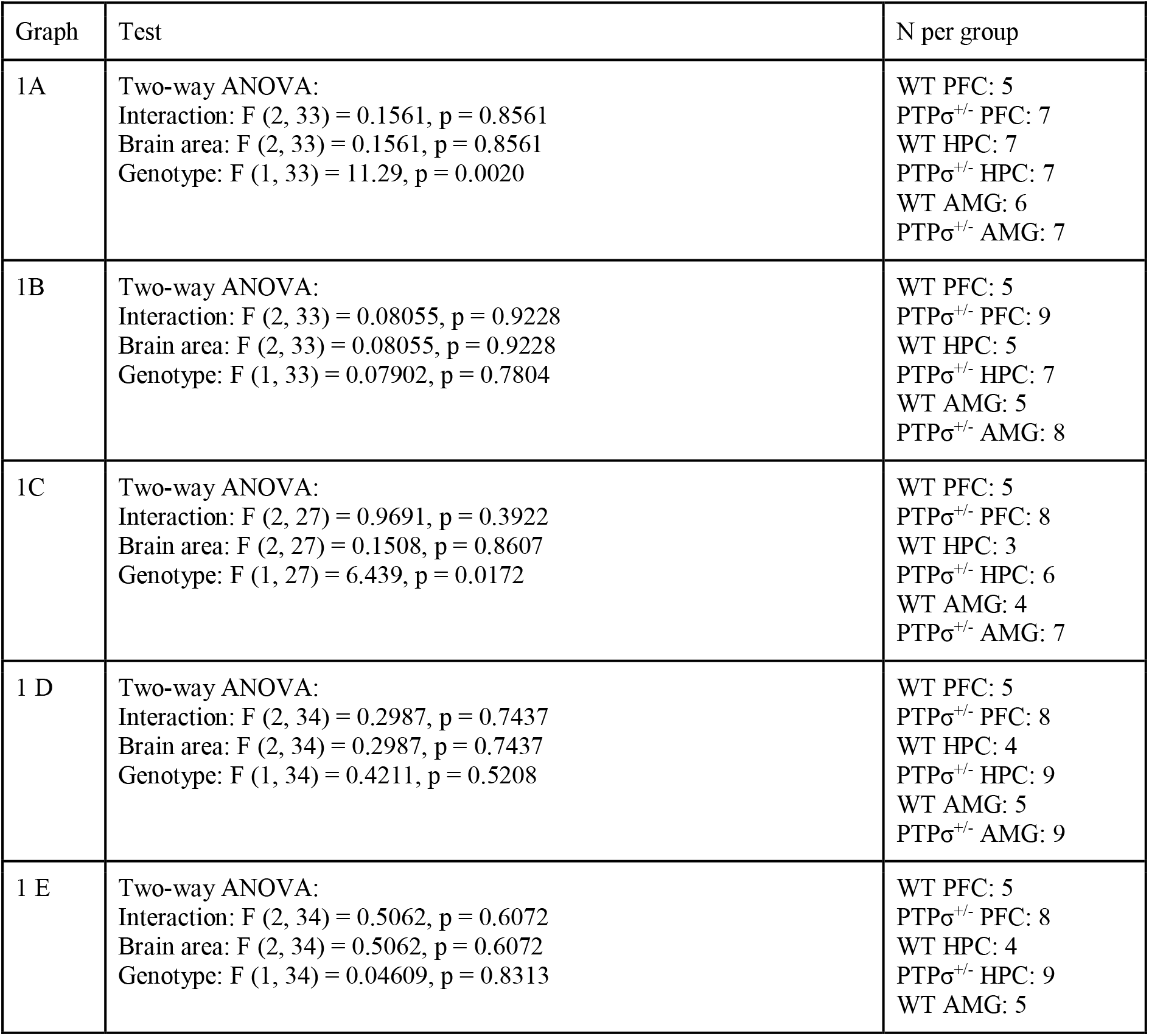
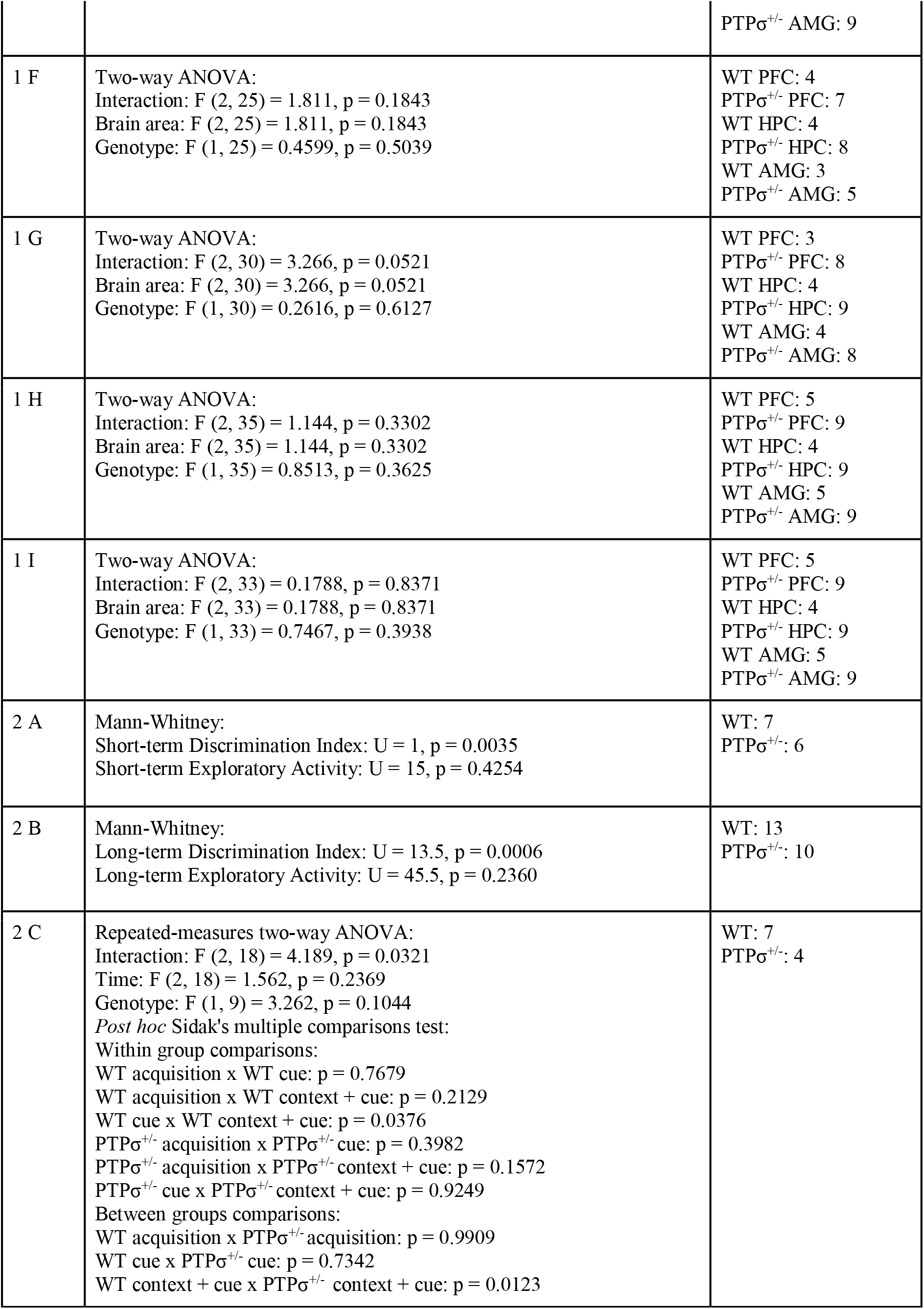
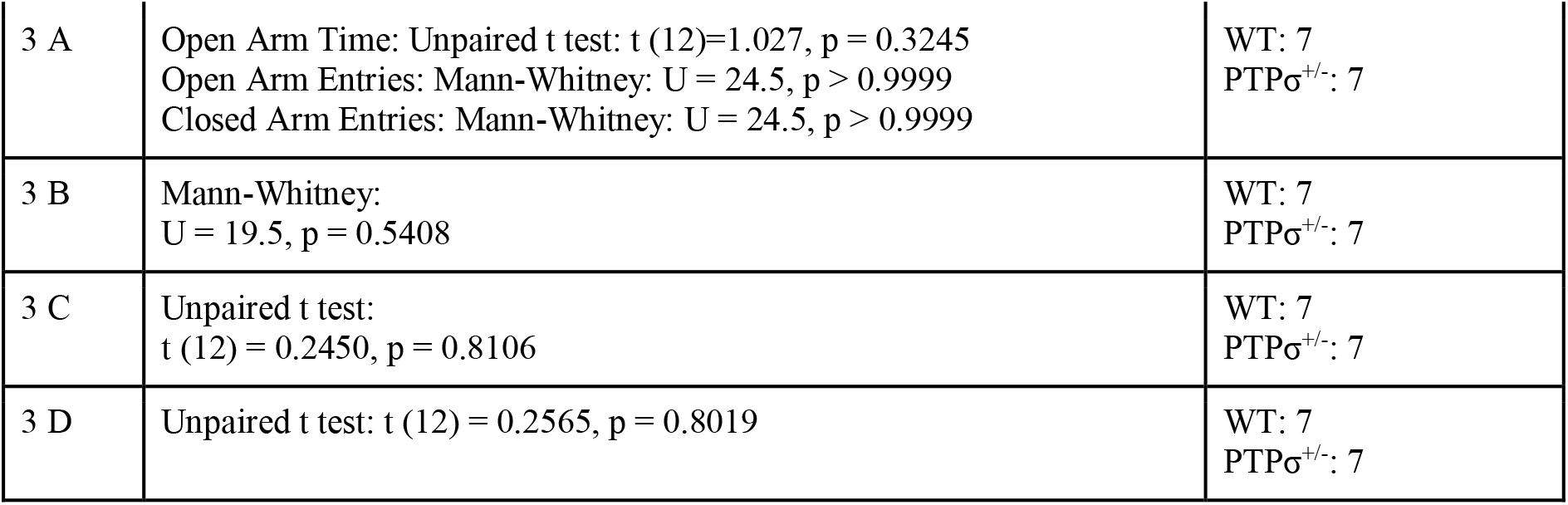
Statistical analysis of the experimental data.

## Results

We have recently demonstrated that PTPσ restricts TRKB signaling *in vitro* and *in vivo* through dephosphorylation, and we have shown that genetic PTPσ deficiency increases basal TRKB phosphorylation and activates network level-plasticity in the visual cortex (Lesnikova et al., 2021). However, it is unknown which intracellular signaling pathways mediate the effects of the increase in pTRKB induced by lack of PTPσ. In order to investigate the molecular profile of changes triggered by PTPσ deficiency, we carried out a biochemical analysis of tissue from different brain areas extracted from PTPσ^+/−^ mice, and compared it with brain tissue from their wild-type littermates.

First, we confirmed that the effect of PTPσ haploinsufficiency on TRKB phosphorylation is not brain region-specific. We checked pTRKB levels in the prefrontal cortex (PFC), hippocampus (HPC) and amygdala (AMG) of PTPσ^+/−^ mice by ELISA, and observed around 50% increase in pTRKB as compared to WT littermates in all of the brain regions tested (Fig. 1A). To confirm that these changes in pTRKB are driven by increased phosphorylation and not by alterations in the TRKB expression, we tested levels of total TRKB protein in the PTPσ^+/−^ mouse samples by Western blot, and detected no changes in total TRKB expression (Fig. 1B). Next, we analyzed phosphorylation levels of phospholipase C gamma 1 (PLCγ1), protein kinase B (Akt), extracellular signal-regulated kinase 1 and 2 (Erk1 and Erk2) and ribosomal protein S6 kinase beta-1 (p70S6 kinase), which are known to be activated downstream of TRKB. We observed a significant increase in pPLCγ1 (Fig. 1C) but not in pAkt, pErk1, pErk2 or p-p70S6 kinase (Fig. 1D, E, F, G correspondingly) in PFC, HPC and AMG of PTPσ^+/−^ mice. Finally, we checked levels of postsynaptic density proteins 93 and 95 (PSD-93 and PSD-95) which are known to be important for synaptic plasticity, but didn’t find any difference in their expression (Fig. 1H and 1I).

**Figure 1.**
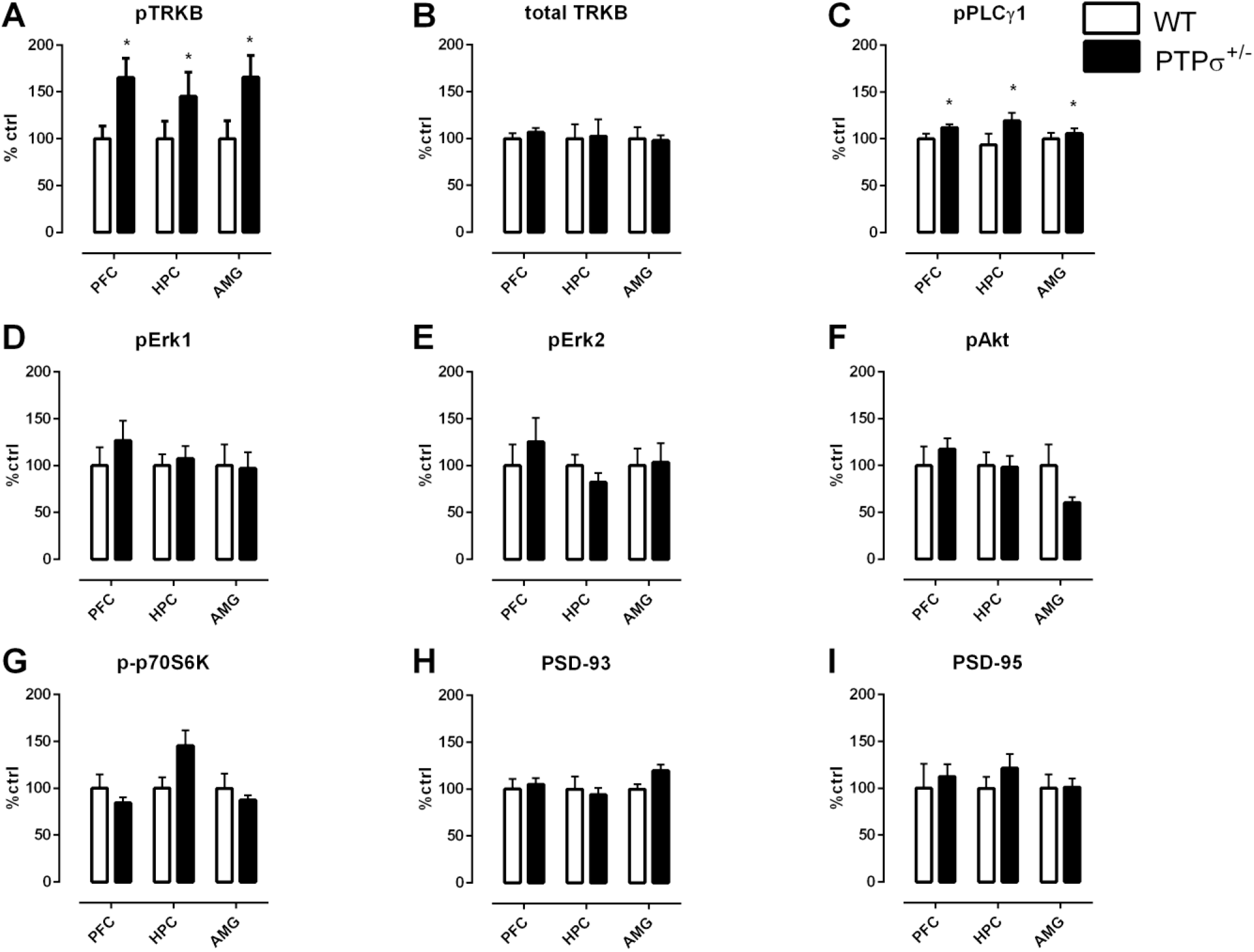
Molecular profile of changes in plasticity-related proteins induced by PTPσ genetic deficiency in different brain areas. **(A)** PTPσ^+/−^ mice have increased phosphorylation of TRKB receptors (pTRKB) and **(B)** no changes in total TRKB levels in the prefrontal cortex (PFC), hippocampus (HPC) and amygdala (AMG). PTPσ^+/−^ mice display **(C)** increased phosphorylation of PLCγ1 (pPLCγ1), no changes in phosphorylation of **(D, E)** Erk, **(F)** Akt, **(G)** p70S6 kinase and no changes in total levels of **(H)** PSD-93 and **(I)** PSD-95 expression. Data on **(A)** was measured by ELISA and on **(B-I)** by Western blot. Data were analyzed by two-way ANOVA. *p < 0.05.

Considering the important role of BDNF-TRKB for various brain functions, including learning and memory (Thoenen, 1995; Minichiello et al., 1999; Cunha et al., 2010), we set out to investigate the behavior phenotype of PTPσ^+/−^ mice. We started with novel object recognition, a cognitive test that evaluates memory-related performance in rodents (Dere et al., 2007). When tested 4 hours after the last training session, PTPσ^+/−^ mice exhibited significantly more interest towards the new object as opposed to the old object, suggesting improved short-term memory (Fig. 2A). This result was similar to the findings by Horn et al., who observed improved short-term recognition memory in full PTPσ knock-out mice (PTPσ^−/−^) (Horn et al., 2012). However, when tested 5 days after the last training session, PTPσ^+/−^ mice demonstrated essentially no difference in exploration between the old and the new object, which indicates that they have significantly impaired long-term recognition memory as opposed to the WT, who remembered the old object and tended to visit the new one more often (Fig. 2B). No difference in total exploratory activity was found between the genotypes in either of the tests.

**Figure 2.**
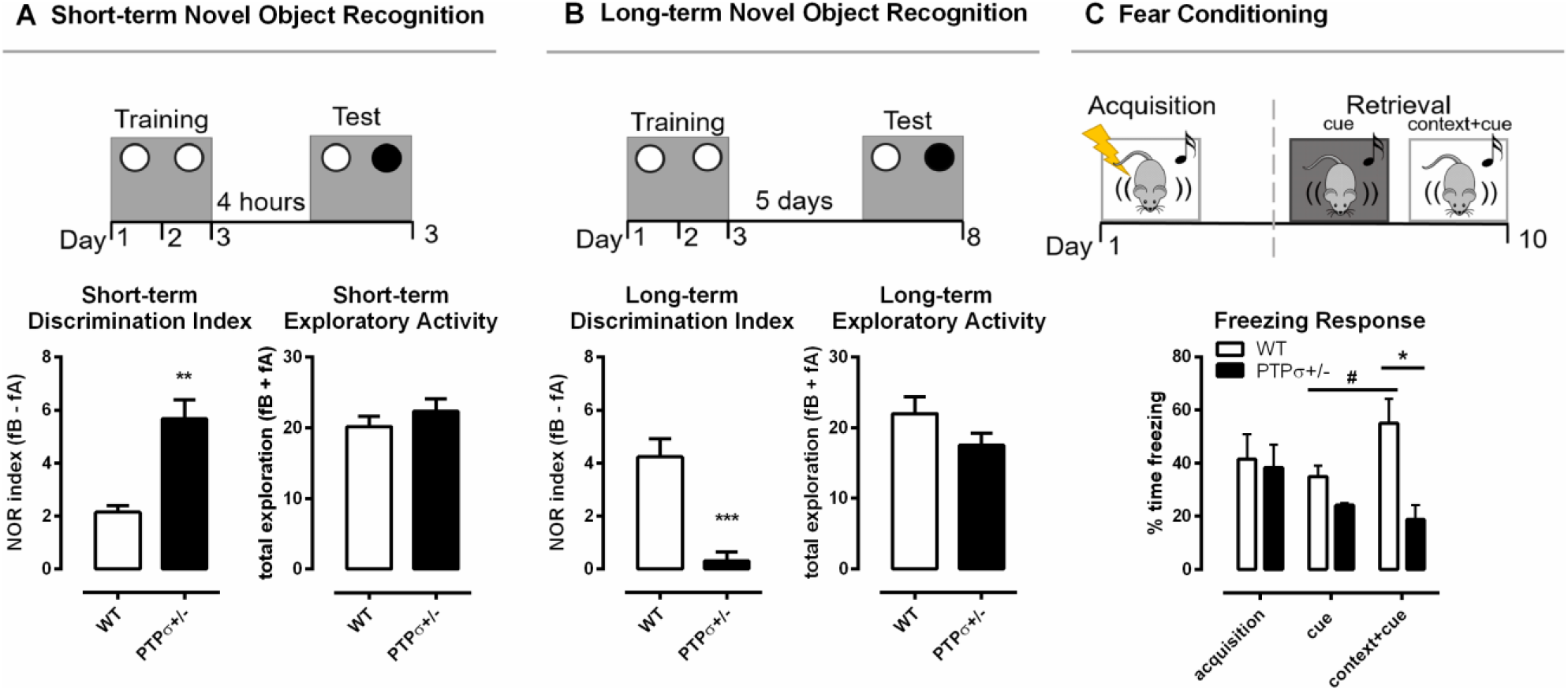
PTPσ^+/−^ mice demonstrate improved short-term but impaired long-term memory. PTPσ^+/−^ mice have **(A)** improved short-term memory and **(B)** impaired long-term memory and no changes in the total exploratory activity in the novel object recognition test. **(C)** Fear response triggered by the cue in the retrieval session was potentiated by the context in WT but not in the PTPσ^+/−^ mice. Data were analyzed by **(A)** and **(B)** Mann-Whitney test, and **(C)** two-way ANOVA.*p < 0.05, **p < 0.01, ***p < 0.001. In 2C: ^#^p < 0.05 (*versus* WT cue), *p < 0.05 (*versus* WT context+cue).

**Figure 3.**
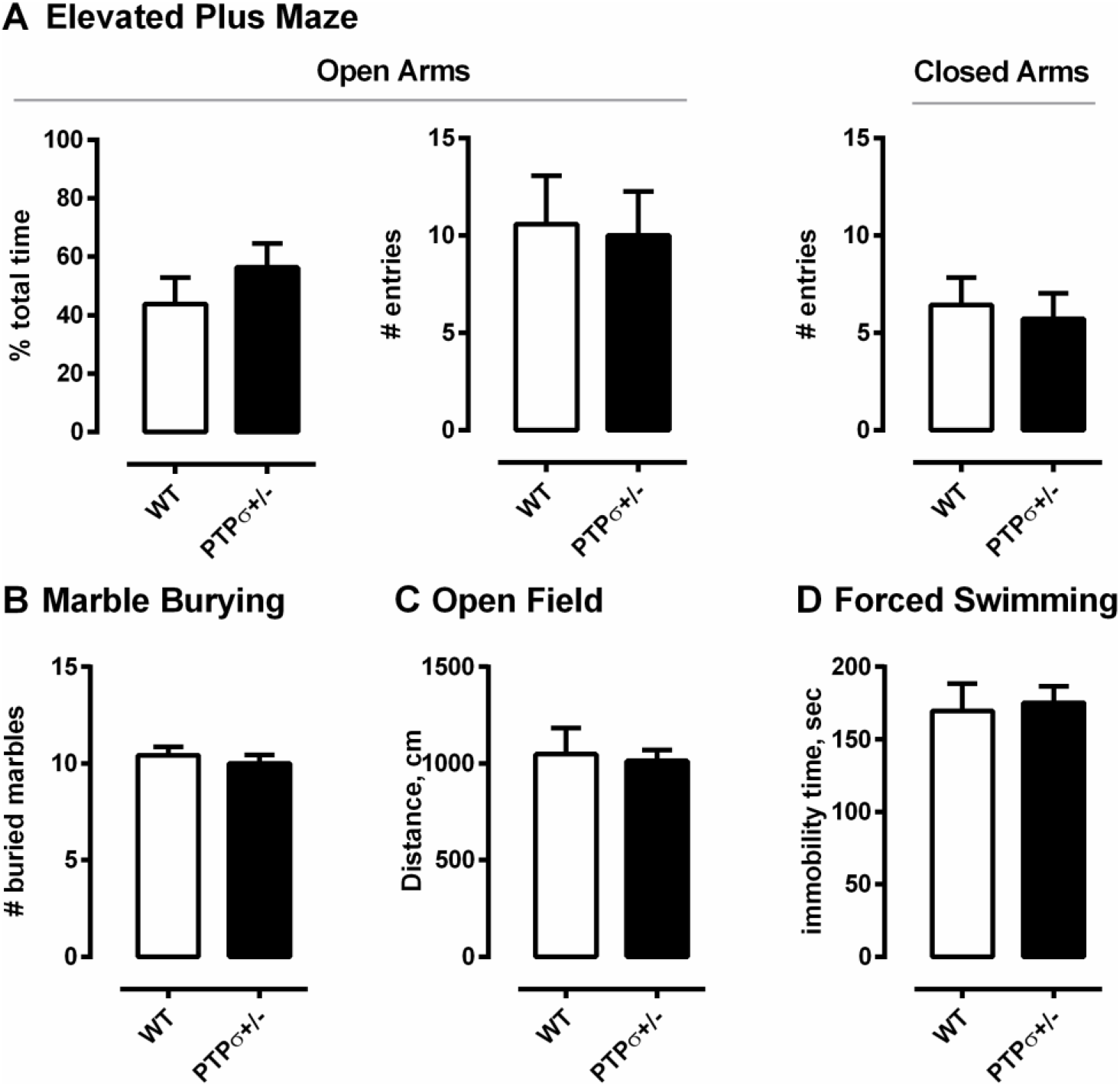
PTPσ^+/−^ mice exhibit normal behavior in tests that do not involve cognitive flexibility. PTPσ^+/−^ mice display no behavioral changes in **(A)** elevated plus maze, **(B)** marble burying, **(C)** open field and **(D)** forced swimming tests as compared to their wild type littermates. Data were analyzed by t-test **(A total time, C, and D)** or by Mann-Whitney test **(A, # entries for Open and Closed Arms, and B)**.

In order to understand if the impaired long-term memory observed in the PTPσ^+/−^ mice was specific to recognition memory or if it was a more generalized memory impairment, we tested those mice also in the fear conditioning model using a protocol that involves both context- and cue-dependent memory components. In the fear conditioning experiment (Fig. 2C), the fear-related memory was established by pairing foot shocks (unconditioned stimulus) with the context (transparent cage with metallic grid floor) and cue (tone). The conditioned response (freezing) was measured immediately after the foot shocks (acquisition) and again 10 days later (retrieval). WT and PTPσ^+/−^ mice displayed similar levels of freezing right after the foot shocks, demonstrating comparable acquisition of fear memories in both genotypes.

Ten days later, we tested memory retrieval triggered by the cue presented either in a novel context (black cage with smooth floor) or in the same context where the foot shocks were previously applied (conditioned context, transparent cage with metallic grid floor). Statistical analysis indicates that there was no significant change in the levels of freezing when acquisition was compared with either of the retrieval sessions, in both genotypes, suggesting that the fear memory did not spontaneously fade away with time. Interestingly, in WT but not in the PTPσ^+/−^ mice, the fear response triggered by the cue in the retrieval session was potentiated by the context, since freezing was greatly increased in the conditioned context in comparison with the novel context in the WT group (indicated by the sharp sign in Fig. 2C). On the other hand, PTPσ^+/−^ mice responded similarly to the cue presented in the novel or in the conditioned context. Finally, the freezing response triggered by the cue combined with conditioned context (cue + context) was significantly higher in the WT compared to the PTPσ^+/−^ mice (indicated by the asterisk in Fig. 2C), reinforcing the evidence that PTPσ deletion might attenuate the retrieval of certain types of long-term memories.

Finally, we ran a behavior battery to test whether PTPσ^+/−^ mice have any behavioral abnormalities other than those related to learning and memory. The tests were separated by at least 24 hours and carried out in the following order: marble burying, elevated plus maze, open field and forced swimming test. Interestingly, PTPσ^+/−^ mice displayed no abnormalities in any of these tests, indicating no difference in the levels of anxiety, total exploratory activity and locomotion as compared to the WT.

## Discussion

We have previously observed that the plasticity-promoting effects of PNN removal are dependent on TRKB signaling in PV interneurons and that PTPσ acts as an agent mediating the opposing effects that perineuronal nets and TRKB have on plasticity (Lesnikova et al., 2021). We demonstrated that PNN-PTPσ-TRKB interaction in parvalbumin interneurons plays a particularly important role, as the chondroitinase ABC effect on network plasticity is abolished in mice lacking full-length TRKB in PV+ cells (PV-TRKB^+/−^ mice) (Lesnikova et al., 2021). Therefore, it is plausible that PV+ interneurons may contribute to the plasticity phenotype of PTPσ mice as they are the major class of neurons expressing PNNs, and are known to be strongly affected by PNN modulation. PV+ cells play a crucial role in many cognitive processes, including learning and memory (Korotkova et al., 2010; Kim et al., 2016), and manipulation of PV+ neurons modulates plasticity states of the neuronal networks (Donato et al., 2013; Winkel et al., 2020). Firing of the PV+ interneurons mediates both memory consolidation (Ognjanovski et al., 2017) and extinction (Trouche et al., 2013; Davis et al., 2017), and optogenetic manipulation of PV+ neurons was sufficient to restore memory-related deficits induced by PNN degradation (Shi et al., 2019). We have shown that optical activation of TRKB in PV+ interneurons is alone sufficient to orchestrate and coordinate large brain networks, rendering them into a state of juvenile-like plasticity (Winkel et al., 2020).

TRKB is a well-characterized positive modulator of plasticity in the brain, essential for learning, memory and other cognitive processes (Yamada and Nabeshima, 2003; Minichiello, 2009). Learning events have been demonstrated to prompt BDNF expression and TRKB phosphorylation (Hall et al., 2000; Gooney et al., 2002), while BDNF mutations in mice and humans are known to induce learning deficits (Linnarsson et al., 1997; Dincheva et al., 2012). TRKB phosphorylation leads to activation of three major downstream signaling pathways: Sch adaptor protein recognizes phosphorylation of tyrosine 515 and activates PI3K/Akt and Ras/MEK/Erk pathways that regulate neuronal survival and neuronal differentiation (Atwal et al., 2000; Minichiello, 2009). PLCγ1 recognizes phosphorylated tyrosine 816 and, once phosphorylated, induces signaling through calmodulin kinase-2 and CREB, thereby modulating neuronal connectivity and plasticity (Patapoutian and Reichardt, 2001; Minichiello, 2009). Specificity of the downstream pathway activation is mediated through different docking partners, and it’s been demonstrated that mutation of PLCγ1 docking site critically affects hippocampal long-term potentiation (LTP), plasticity and learning (Minichiello et al., 2002; Gruart et al., 2007). Surprisingly, a similar mutation of the Shc recognition site, which mediates Akt and Erk activation, produces only minimal effects on BDNF signaling *in vivo* (Minichiello et al., 1998). Given that PTPσ^+/−^ mice demonstrate an increase in PLCγ1 activation, their phenotype of facilitated plasticity and learning is not entirely surprising. However, the reason for specificity of activation for this particular pathway remains to be further investigated.

TRKB overexpression in a transgenic mouse line (TRKB.TK+) has yielded similar results with positive modulation of PLCγ1 but not PI3K/Akt or Erk pathways (Koponen et al., 2004). These TRKB.TK+ mice demonstrated facilitated learning (tested in Morris water maze) similar to PTPσ^+/−^ mice. However, TRKB.TK+ mice also showed improved long-term memory in both Morris water maze and fear conditioning paradigms (Koponen et al., 2004), as opposed to PTPσ^+/−^ mice having impaired long-term fear conditioning and object recognition memory. However, direct comparison of these observations is difficult due to the timeline differences between the tests and because TRKB.TK+ mice overexpress TRKB in pyramidal cells and not in PV neurons. TRKB.TK+ were tested 48 hours after training at the latest, when the memory traces were relatively fresh, while PTPσ^+/−^ mouse memory was assessed 5-10 days after the initial learning. Moreover, even though BDNF-TRKB signalling is known to be critical for both short-term and long-term memory function (Bekinschtein et al., 2014), excessive BDNF-TRKB activation can also render previously established long-term memories labile and promote their erasure. For example, direct BDNF infusion into infralimbic prefrontal cortex facilitates extinction of fear and drug-induced conditioned place preference memories (Peters et al., 2010; Otis et al., 2014; Kataoka et al., 2019), with similar effects achieved by TRKB facilitation by chronic antidepressant treatment (Karpova et al., 2011). Even though time and region-specific mechanisms of BDNF-TRKB role in memory events remain to be fully clarified, it is plausible to assume that TRKB activation has to be time-precise and transient in order to lead to robust memory acquisition. Since PTPσ^+/−^ mice have chronically increased TRKB activation due to reduced dephosphorylation from PTPσ, their neuronal networks seem to acquire a state of hyperplasticity, or tonic plasticity (as opposed to the phasic in the normal brain), which allows for enhanced acquisition of memories but not for their retention.

It is interesting that a similar dissociation between short- and long-term memories can be experimentally induced by PNN manipulation. PNN downregulation by chABC digestion or genetic knockdown of their components has been shown to improve object recognition memory for a period up to 48 hours (Romberg et al., 2013; Rowlands et al., 2018). However, perineuronal nets are also indispensable for long-term memory retention, as PNN degradation has been demonstrated to impair persistence of memories and promote their erasure in different brain areas such as prefrontal cortex (Hylin et al., 2013; Slaker et al., 2015), hippocampus (Hylin et al., 2013; Riga et al., 2017; Shi et al., 2019) and amygdala (Gogolla et al., 2009; Xue et al., 2014; Pignataro et al., 2017). Since PTPσ^+/−^ mice have a lower number of PNN receptors available for transduction of their signals, the PNN network consolidation function in their brain is likely to be compromised, which may improve short-term but impair long-term memory performance.

PTPσ is a complex molecule attributed to a number of different functions. Apart from being a phosphatase and a receptor for PNNs, it is also known to be a synaptic adhesion molecule, playing an important role in excitatory synapse assembly. It binds a number of partners in both cis- and trans-synaptic manner, including SLIT and NTRK-like proteins (Slitrks) (Um et al., 2014), synaptic adhesion-like molecule 3 (SALM3) (Li et al., 2015b), leucine-rich repeat transmembrane protein 4 (LRRTM4) (Ji et al., 2015), liprin-α (Bomkamp et al., 2019) and tropomyosin receptor kinase C (TRKC) (Takahashi et al., 2011; Coles et al., 2014). These interactions have been demonstrated to be critical for normal excitatory synaptic formation. According to the model proposed in the literature, PTPσ drives excitatory synapse formation through a combination of trans-synaptic interactions with partners such as TRKC (but not TRKB) and Slitrks, and intracellular signaling cascades when PTPσ intracellular domains interact in cis with adaptor proteins (such as liprin-α) and direct substrates, such as N-cadherin or TRKB (Coles et al., 2011; Han et al., 2018; Bomkamp et al., 2019). Global PTPσ knockout and conditional deletion of PTPσ impairs excitatory synapse transmission, leading to abnormalities in synaptic structure (Horn et al., 2012; Han et al., 2020) and reduced number of excitatory synapses (Han et al., 2018, 2020). PTPσ deficiency also increases frequency but reduces efficiency of excitatory postsynaptic currents (Horn et al., 2012; Han et al., 2020) and impairs long-term potentiation (LTP) (Horn et al., 2012). LTP deficits were also observed when PTPσ knockdown was restricted to excitatory cortical and hippocampal neurons (Kim et al., 2020). Similarly, overexpression TRKB in pyramidal neurons, which increases TRKB phosphorylation and signaling through PLCγ1, also impairs LTP (Koponen et al., 2004). Interestingly, Kim et al. showed that presynaptic PTPσ deficiency selectively impairs NMDA-dependent postsynaptic currents, and it happens through a mechanism independent of PTPσ trans-synaptic adhesion, suggesting further involvement of PTPσ cytoplasmic phosphatase domains in this process (Kim et al., 2020). Even though direct relationship between LTP and memory remains to be a subject to debate (Stevens, 1998; Nicoll, 2017), NMDA-dependent LTP is known to be critical for normal memory performance, and disturbances in this pathway impair memory function (Nakazawa et al., 2004; Li and Tsien, 2009; Morris, 2013; Nabavi et al., 2014). Since PTPσ plays a substantial role in excitatory synaptic transmission and normal long-term potentiation, its partial deletion from PTPσ^+/−^ mouse brain may cause deficiencies in normal synapse function and stabilization, leading to an inability of the synapses to efficiently support long-term memory maintenance.

Evanescence of long-term memories may be a disadvantage; however, it may also be beneficial under certain conditions when memories carry over a trauma such as in PTSD. Labilization of memories has been proposed to be a potential therapeutic tool for such disorders (Kindt et al., 2009), and treatment strategies targeting both PNNs (Gogolla et al., 2009; Banerjee et al., 2017; Pignataro et al., 2017) and BDNF-TRKB system (Karpova et al., 2011; Andero and Ressler, 2012; Kataoka et al., 2019) were suggested to carry translational promise. Now that PTPσ has been established as a missing link in the PNN-PTPσ-TRKB axis in PV interneurons, its further investigation as a potential target for PTSD treatment is promising. Moreover, intracellular sigma peptides (ISP) that bind to PTPσ intracellular domain and inhibit its phosphatase activity have recently been developed and successfully tested in the rodent models of spinal cord injury (Li et al., 2015a) and multiple sclerosis (Luo et al., 2018; Tran et al., 2018). ISPs are currently undergoing an FDA approval procedure and are being prepared for the first clinical trials, which bears promise for their future use in patients. However, their further development and application in humans must be carried out with caution, as long-term memory loss may come up as a potential side-effect of ISP treatment.

## Acknowledgments

We thank Nelli Koivisto and Vootele Võikar (Mouse Behavioral Phenotyping Facility, Laboratory Animal Center - University of Helsinki) for their expert advice on animal research and Sulo Kolehmainen and Seija Lågas for technical help.

